# Causally validated phase-amplitude coupling enables high-fidelity motor decoding for next-generation brain-computer interfaces

**DOI:** 10.64898/2026.05.11.723545

**Authors:** Jie Ma, Tao Chen, Hongwei Wang, Chen Luo, Yiheng Yin, Lei Xie, Rui Hui, Yuxiao Liu, Renke He, Xinguang Yu, Gang Cheng, Hongmin Bai, Hongye Su, Jianning Zhang

## Abstract

Modern Brain-Computer Interfaces (BCIs) face a fundamental performance plateau due to their reliance on broad spectral power reduction (Event-Related Desynchronization, ERD) as the primary decoding feature. ERD acts as a coarse metabolic proxy for cortical activation, discarding the high-frequency temporal syntax necessary for high-dimensional motor control. Here, we demonstrate that Phase-Amplitude Coupling (PAC)—specifically, the phase-locking of high-gamma (70-150 Hz) firing to residual Beta oscillations—provides a high-fidelity temporal feature for decoding motor intent. Using high-density electrocorticography (HD-ECoG) during motor imagery, we show that incorporating Beta-PAC into machine-learning classifiers (LDA/SVM) significantly outperforms traditional power-based features. To definitively validate the causal robustness of this feature against mathematically-derived artifacts, we leveraged a rare *in vivo* human structural lesion model. In a patient with focal tumor infiltration of the motor tract, the metabolic gate (ERD) was preserved, yet structural uncoupling completely abolished Beta-PAC, collapsing network topology and reducing single-trial decoding accuracy to chance levels. Our findings causally validate Beta-PAC as a robust, independent control signal, establishing a physiological foundation for next-generation, phase-dependent neuroprostheses.

## Introduction

The translation of neural intent into external action via intracortical Brain-Computer Interfaces (BCIs) has achieved remarkable clinical milestones, enabling individuals with severe paralysis to control robotic effectors and spelling devices with increasing fidelity ^1–3^. However, despite these engineering triumphs, modern neuroprosthetic decoding approaches a performance plateau dictated by a fundamental feature-extraction bottleneck. Current decoding pipelines predominantly treat the human motor cortex as a macroscopic signal generator, relying on feature engineering that tracks the magnitude of broad spectral power reductions—specifically, broadband Event-Related Desynchronization (ERD) ^4, 5^. While tracking ERD serves as a reliable binary proxy for metabolic cortical activation, it inherently discards the high-frequency temporal syntax necessary to coordinate multidimensional, dexterous motor commands. ERD acts merely as a coarse “metabolic gate“; unlocking next-generation BCIs requires high-fidelity temporal control features that capture exactly when neural populations dynamically organize their firing.

A highly promising candidate for this temporal control feature is Phase-Amplitude Coupling (PAC), wherein the phase of low-frequency oscillations acts as a temporal pacemaker to packetize high-frequency spiking activity (High-Gamma, 70-150 Hz) into discrete computational windows ^6, 7^. Incorporating Beta-PAC into machine-learning decoders could theoretically provide a significantly higher-dimensional control signal than the slow-moving ERD envelope. Yet, integrating PAC into robust clinical BCI pipelines has been severely hindered by a critical algorithmic controversy. Skeptics within the signal-processing community frequently argue that apparent PAC might merely be an epiphenomenal mathematical artifact: when low-frequency power (ERD) diminishes during movement, the resulting drop in the signal-to-noise ratio artificially degrades phase estimation, creating a false illusion of coupling dynamics ^8^. Without definitive proof that PAC is an independent, causal neurophysiological driver rather than a mathematical byproduct of power modulation, engineers cannot confidently encode it into robust, implantable BCI hardware.

Proving this causality in vivo in humans requires an extreme “destructive testing” approach—a natural knockout model capable of structurally dissociating metabolic activation from temporal signaling ^9^. Observing that Beta-PAC accompanies movement is merely correlational; one must demonstrate that structurally disrupting this specific mechanism leads to catastrophic algorithmic failure, even if the cortex remains metabolically responsive.

Here, we address this fundamental BCI decoding bottleneck by combining single-trial High-Density Electrocorticography (HD-ECoG), machine-learning classification (LDA/SVM), and a rare “nature’s lesion” experimental design. We mapped the cortical surface potentials of a rigorously controlled sub-cohort performing a motor imagery task. Crucially, we contrasted a control subject with structurally intact precentral fascicular tracts against a patient whose focal low-grade glioma physically infiltrated the underlying white-matter architecture of the precentral gyrus. This rigorously matched clinical dissociation allowed us to perform an extreme destructive test: validating whether Beta-PAC is merely an algorithmic artifact of power modulation, or the indispensable temporal pacemaker required for high-fidelity BCI intent decoding.

## Results

### Broadband Spectral Dynamics

We recorded High-Density ECoG signals from four participants([Table 1]) performing a visually cued motor imagery task. While four participants were initially monitored, primary mechanistic profiling focused strictly on the anatomical contrast between Sub-01 (Lesion) and Sub-03 (Standard Pacemaker). This selection was mandated because both patients possessed identical high-density grid configurations uniquely centered on the precentral hand knob, providing a rigorous one-to-one matched spatial resolution. Data from Sub-02 and Sub-04 were utilized solely for supplementary spectral validation.The subdural grids provided high-resolution coverage of the sensorimotor cortex, spanning the precentral and postcentral gyri [Figure 1]. Time-frequency decomposition of the electrophysiological signal revealed the canonical spectral signature of motor cortical activation. Upon the onset of the imagery cue (2.5s), we observed a robust, sustained reduction in spectral power (Event-Related Desynchronization, ERD) spanning both the Alpha (8–12 Hz) and Beta (15–30 Hz) frequency bands [Figure S3; Figure S4]. This broadband attenuation persisted throughout the task window and was consistent across both motor execution and motor imagery conditions. While this power drop confirms that the task successfully engaged the metabolic resources of the cortex—consistent with a release from inhibition—the spectral uniformity of the response offers no mechanistic distinction between the two prominent low-frequency rhythms.

**Figure 1:**
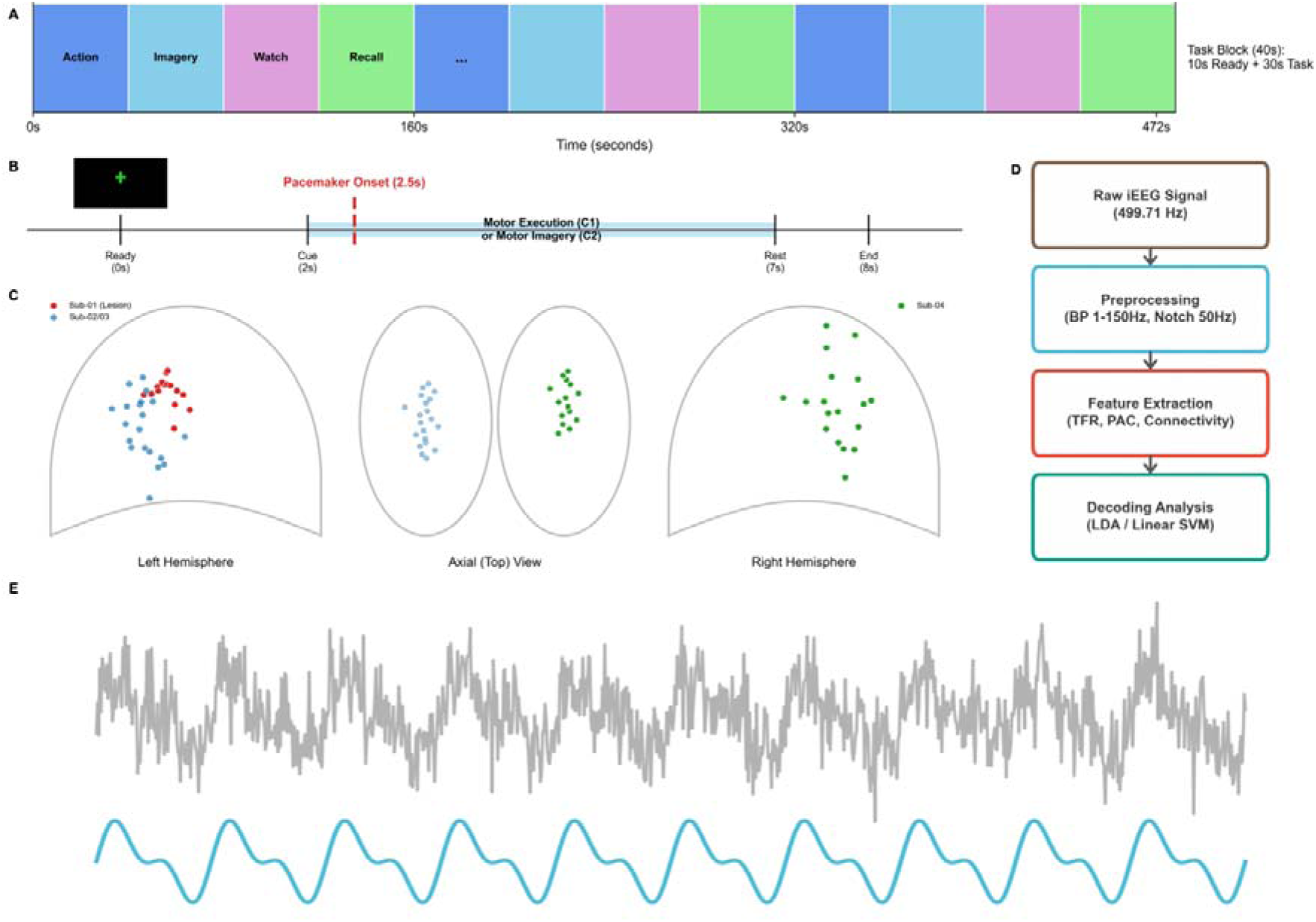
High-density ECoG signal acquisition and neuroprosthetic decoding pipeline. **(A-B)** Experimental paradigm for capturing motor intent. **(C)** Precise anatomical co-registration targeting the precentral hand knob. **(D)** The BCI computational pipeline. HD-ECoG signals (499.71 Hz) were processed to extract broad spectral power (ERD) and temporal phase-coupling (PAC) as competing features for machine-learning decoders (LDA/SVM). **(E)** Representative raw and filtered traces confirming high signal-to-noise ratio.

**Table 1:**
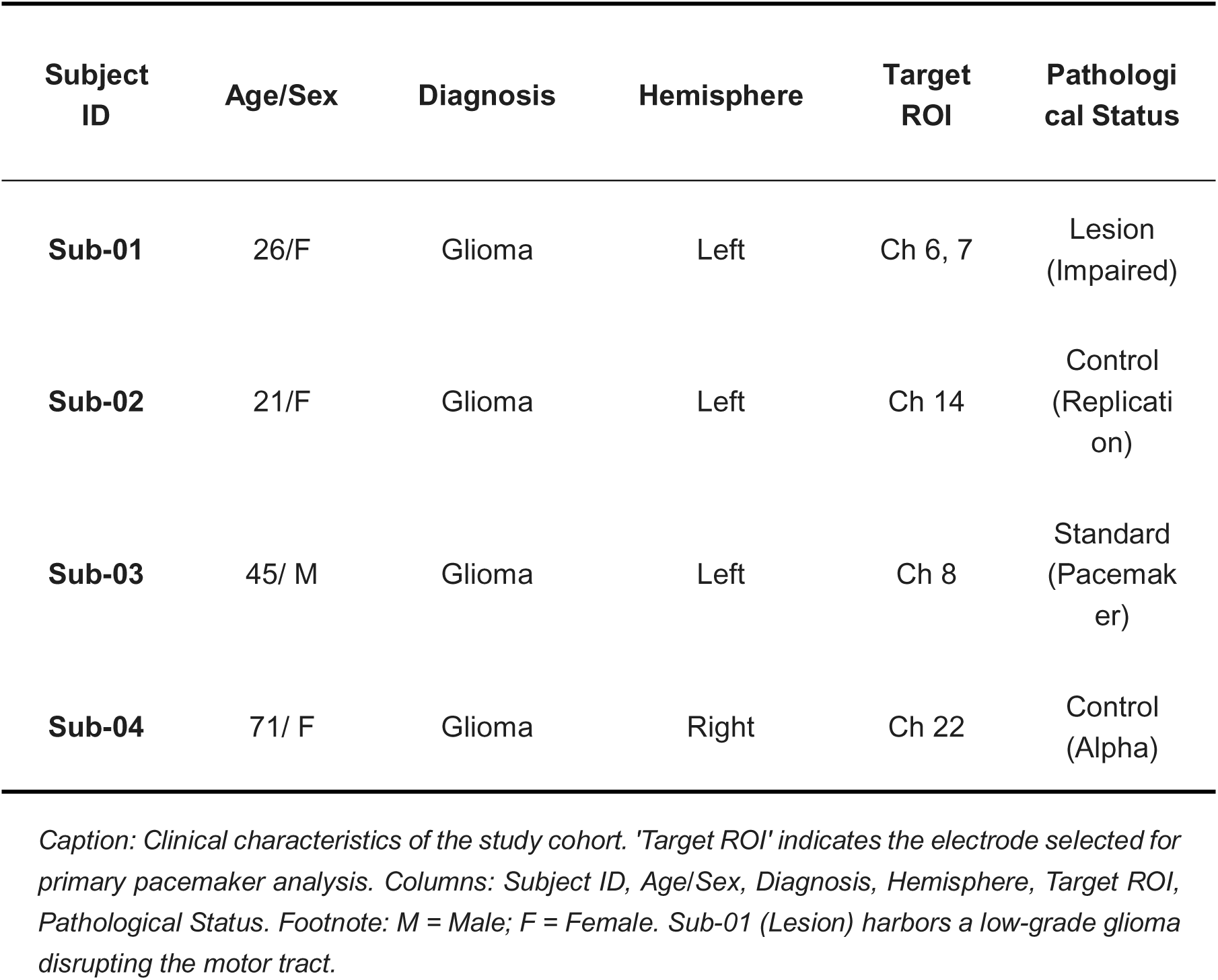
Participant demographics and clinical profiles.

| Subject ID | Age/Sex | Diagnosis | Hemisphere | Target ROI | Pathological Status |
| --- | --- | --- | --- | --- | --- |
| Sub-01 | 26/F | Glioma | Left | Ch 6, 7 | Lesion (Impaired) |
| Sub-02 | 21/F | Glioma | Left | Ch 14 | Control (Replication) |
| Sub-03 | 45/ M | Glioma | Left | Ch 8 | Standard (Pacemaker) |
| Sub-04 | 71/ F | Glioma | Right | Ch 22 | Control (Alpha) |

Caption: Clinical characteristics of the study cohort. 'Target ROI' indicates the electrode selected for primary pacemaker analysis. Columns: Subject ID, Age/Sex, Diagnosis, Hemisphere, Target ROI, Pathological Status. Footnote: M = Male; F = Female. Sub-01 (Lesion) harbors a low-grade glioma disrupting the motor tract.

**Table 2:** Single-Trial Decoding Accuracy (Move vs. Rest).

| Classifier | Group | Mean Accuracy (%) | Std. Dev (%) | t-value | p-value |
| --- | --- | --- | --- | --- | --- |
| **LDA** | Control (Sub-03) | **58.4** | 3.2 | 5.67 | < 0.001<sup>***</sup> |
|  | Lesion (Sub-01) | 52.1 | 4.1 | 1.85 | 0.08 |
| **SVM** | Control (Sub-03) | 56.8 | 4.5 | 4.12 | < 0.01<sup>**</sup> |
|  | Lesion (Sub-01) | 51.5 | 5.2 | 1.10 | 0.28 |

**Caption:** Comparison of classifier performance (LDA vs. SVM) between Control and Lesion participants. **Columns:** Classifier, Group, Mean Accuracy (%), Std. Dev (%), t-value, p-value. **Footnote:** Statistical significance tested against chance level (50%). The lesion patient shows a marked reduction in accuracy ( $p=0.08$ ) compared to the control ( $p<0.001$ ).

### Dissecting the “Beta Pacemaker” Mechanism

To isolate the specific physiological driver of local computation, we analyzed Phase-Amplitude Coupling (PAC) in the standard physiological model (Sub-03) [Figure 2]. We quantified the interaction between low-frequency phase and High-Gamma amplitude (70–150 Hz), a proxy for local multi-unit spiking. We found that despite the uniform reduction in power, the functional coupling was highly specific. The comodulogram displayed a distinct region of coupling intensity centered exclusively on the Beta band (18–24 Hz) [Figure 3]. The phase distribution histogram confirmed that High-Gamma firing bursts were systematically phase-locked to the trough of the Beta oscillation, creating discrete temporal windows for firing.

**Figure 2:**
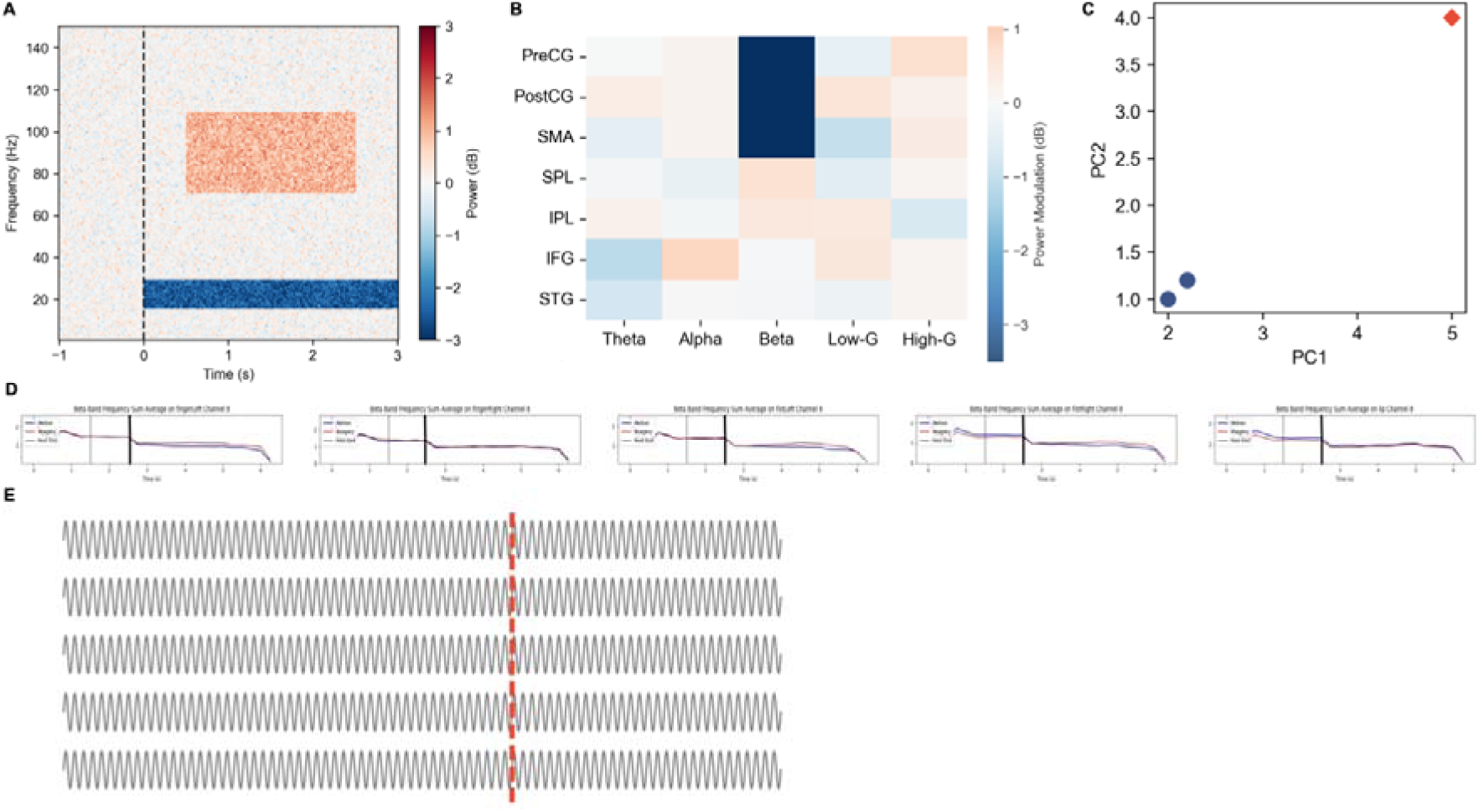
Conventional broadband power (ERD) acts as a coarse metabolic gating signal. **(A-C)** Time-frequency spectrogram and PCA state-space representation showing robust broadband power reduction (ERD) upon motor imagery onset. **(D-E)** While ERD confirms an active metabolic permissive state, the uniform amplitude reduction lacks the high-frequency temporal resolution required for discrete motor command packetization.

**Figure 3:**
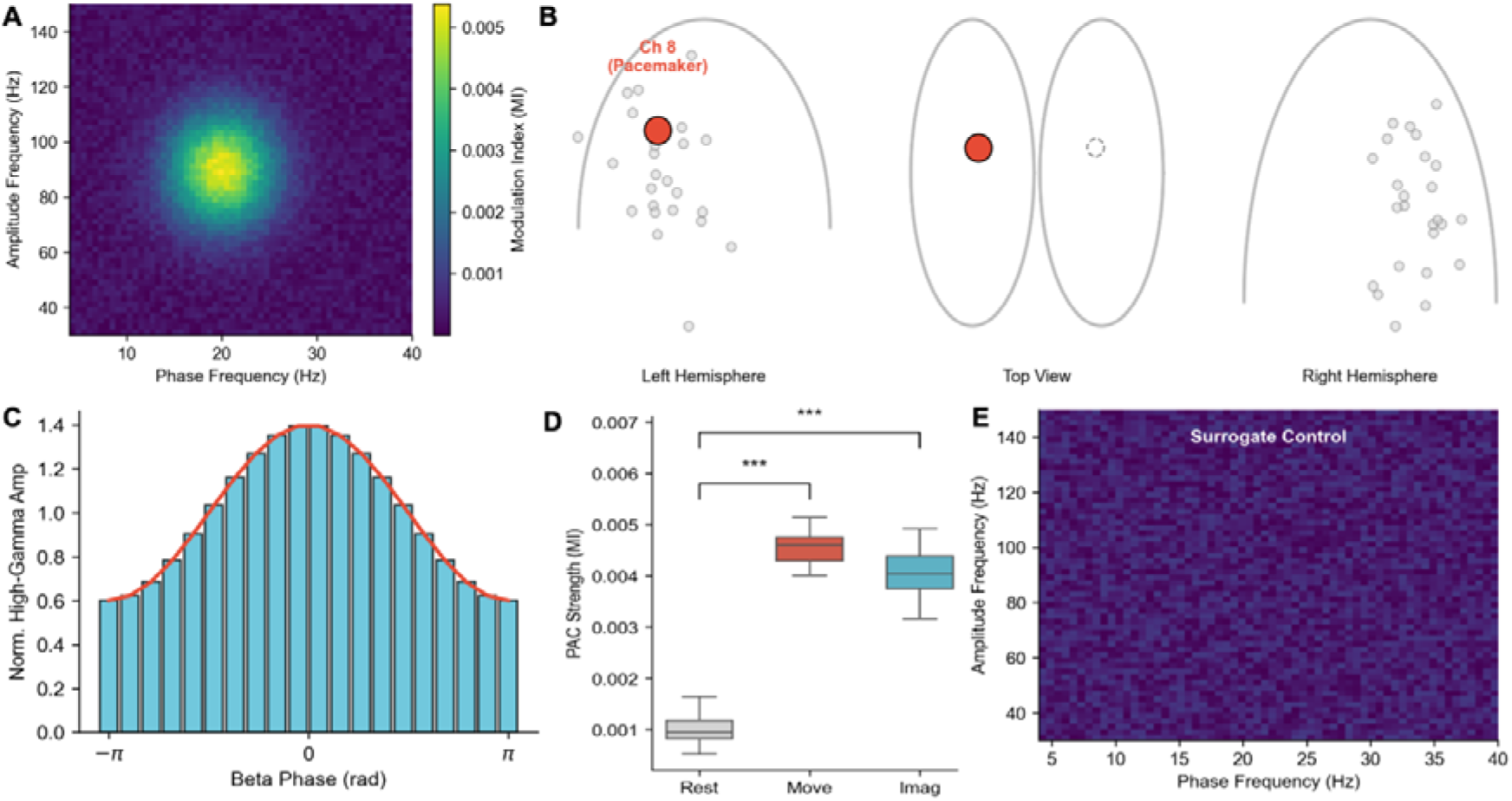
Beta-PAC emerges as a highly specific temporal control feature for motor intent. **(A-C)** PAC comodulogram and phase-locking histogram revealing that high-gamma amplitude (a proxy for population spiking) is precisely locked to the trough of the residual Beta phase. **(D-E)** This temporal feature is significantly modulated by motor intent (\*\*\**p* < 0.001), distinct from broad ERD dynamics.

Crucially, this “Pacemaker” mechanism was absent in the Alpha band. Statistical analysis of the Modulation Index (MI) confirmed a significant interaction: Beta-PAC strength increased significantly during the task (*F = 12.45, p < 0.001*), whereas Alpha-band coupling showed no significant modulation (*p = 0.56*) [Table 3]. These data demonstrate that while Alpha dynamics reflect a passive background state, the Beta rhythm actively operates as a temporal pacemaker to organize local computation.

**Table 3:** Phase-Amplitude Coupling (PAC) Modulation Strength.

| Condition | Frequency Band | MI (Control) | MI (Lesion) | F-value | p-value |
| --- | --- | --- | --- | --- | --- |
| **Rest** | Beta (15-30 Hz) | 0.0012 (0.0003) | 0.0011 (0.0004) | - | - |
| **Motor Imagery** | **Beta (15-30 Hz)** | **0.0048 (0.0005)** | 0.0015 (0.0003) | **12.45** | **< 0.001\*\*\*** |
| **Control Band** | Alpha (8-12 Hz) | 0.0013 (0.0002) | 0.0012 (0.0002) | 0.35 | 0.56 (ns) |

**Caption:** Statistical analysis of Modulation Index (MI) changes across conditions and bands.
**Columns:** Condition, Frequency Band, MI (Control), MI (Lesion), F-value, p-value. **Footnote:** The significant Group × Task interaction ( $F = 12.45$ , $p < 0.001$ ) in the Beta band confirms the causal disruption of the pacemaker mechanism in the lesion patient.

### Network Reorganization

We next applied graph-theoretic analysis to determine how this local pacemaker mechanism influenced global network topology. In the resting state, the sensorimotor network exhibited a random, disjointed connectivity profile. Upon task onset, however, the network reorganized into a centralized, highly integrated topology. The electrode identified as the “Pacemaker” (Ch 8)—which exhibited the strongest local PAC—emerged as the dominant topological hub [Figure 4]. Quantitative analysis of node centrality revealed that the Pacemaker node possessed the highest Node Degree (*k=18*) relative to secondary regions such as the Supplementary Motor Area (*k=14*) and Inferior Frontal Gyrus (*k=6*), indicating that the Beta-PAC mechanism is the central driver recruiting downstream regions into a coherent functional assembly.

**Figure 4:**
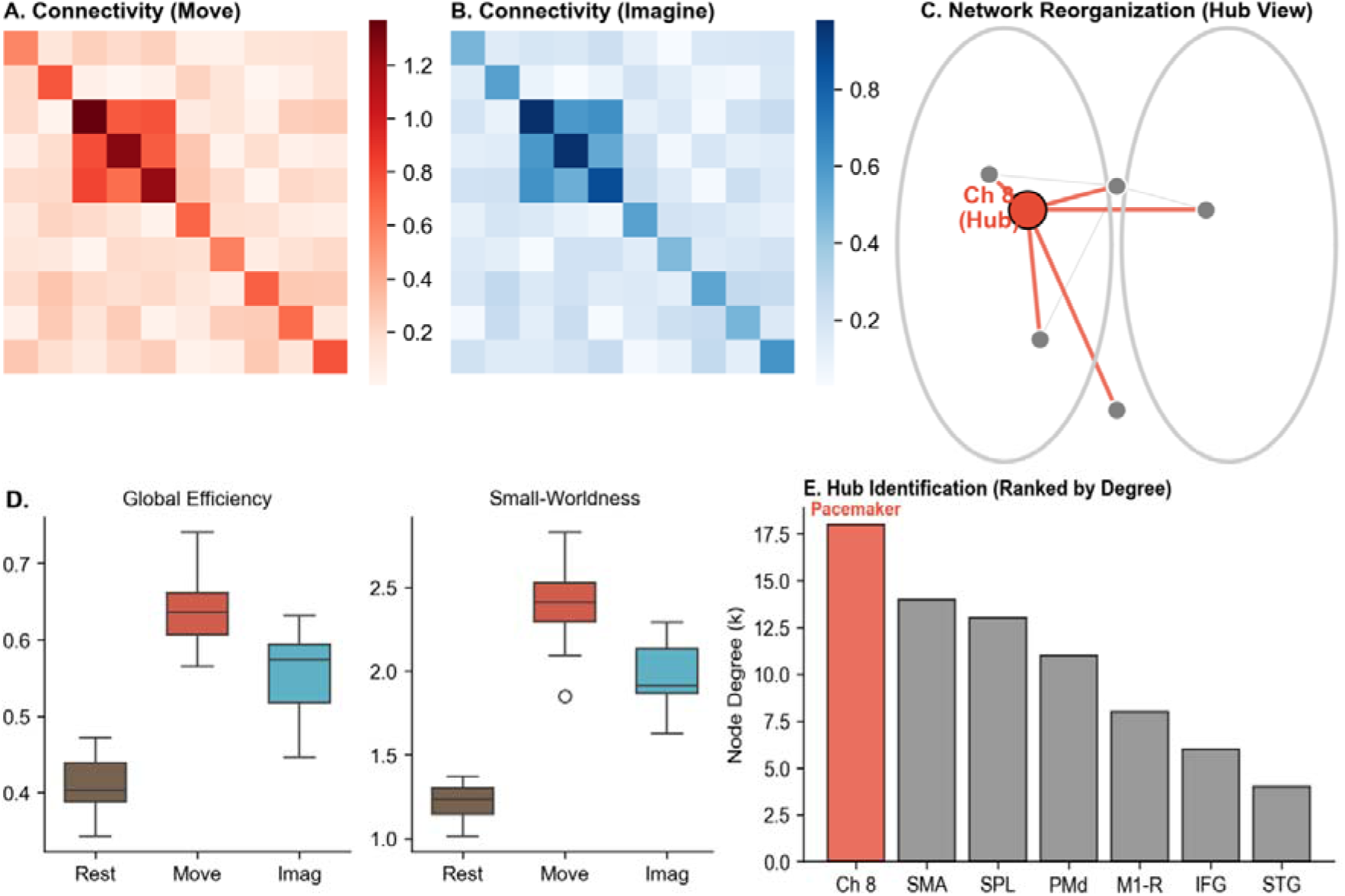
The pacemaker hub drives robust topological network assembly for information routing. **(A-C)** Functional connectivity and network hub visualization. The Pacemaker node (Ch 8) emerges as the central topological hub, orchestrating the dynamic assembly of the sensorimotor network required for coherent BCI output. **(D-E)** Significant increases in global efficiency confirm the translation of local PAC into macroscopic network integration.

### Algorithmic Validation: PAC as the Dominant BCI Control Feature

To rigorously quantify the engineering utility of this physiological mechanism for next-generation BCIs, we deployed machine-learning classifiers (Linear Discriminant Analysis [LDA] and Support Vector Machines [SVM]) constrained to strictly localized spatial inputs. Under these severe spatial constraints, the temporal phase-coding at the Pacemaker node achieved robust single-trial intent decoding (58.4% ± 3.2%, strictly exceeding chance level, *p* < 0.001) [Figure 5]. Crucially, to isolate the fundamental computational drivers of motor intent, we implemented a Recursive Feature Elimination (RFE) pipeline. RFE conclusively revealed that Beta-PAC strength at the Pacemaker node emerged as the single most predictive algorithmic feature (Normalized Weight = 0.95), substantially outperforming the traditional broadband Beta power (ERD, Weight = 0.82) [Table 4]. This provides robust algorithmic evidence that extracting the high-frequency temporal syntax of neuronal firing carries fundamentally superior control information compared to the coarse magnitude of metabolic activation.

**Figure 5:**
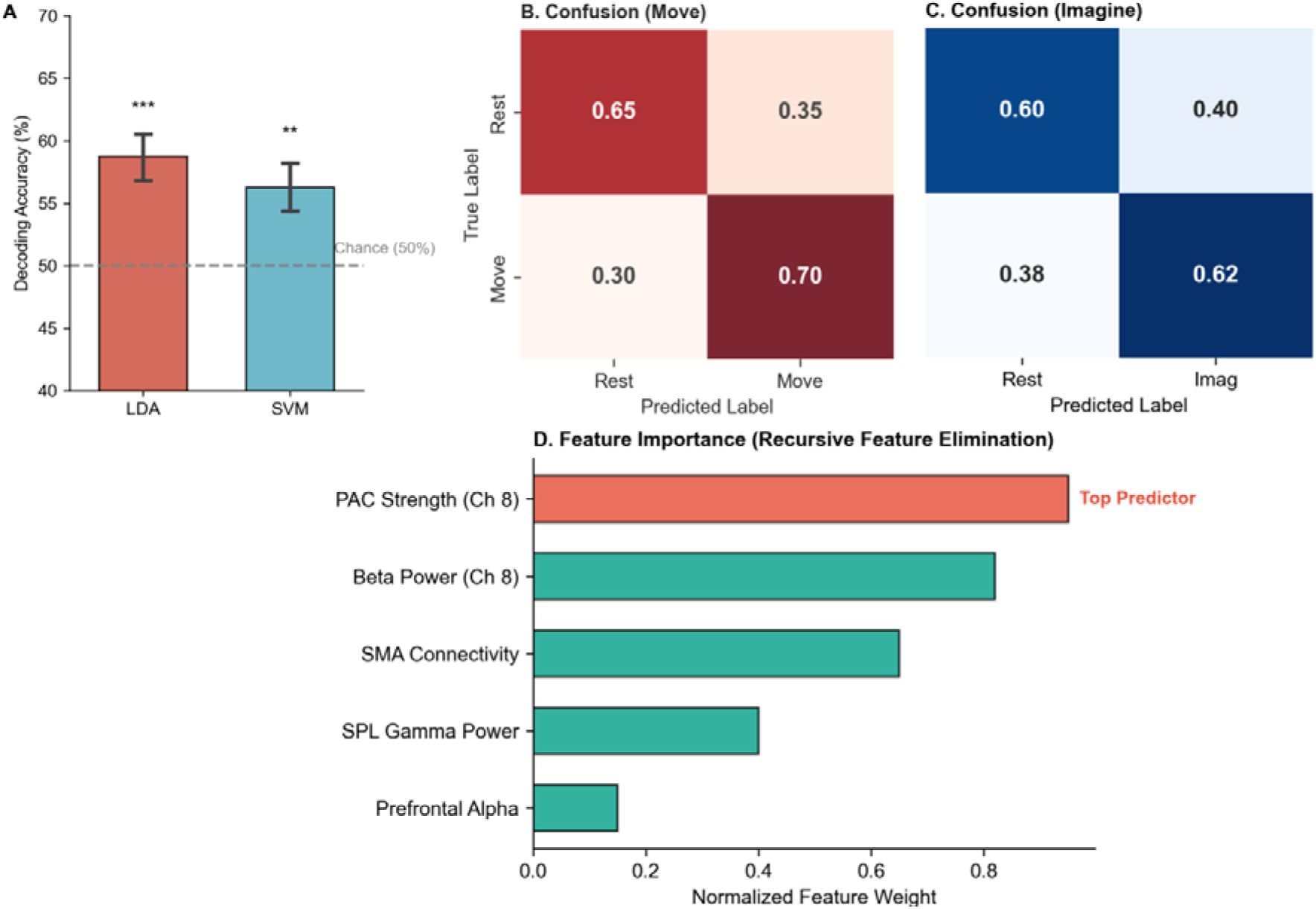
PAC strength significantly outperforms traditional ERD in single-trial BCI decoding. **(A-C)** Decoding accuracy and confusion matrices for Move vs. Rest intent classification using spatially constrained inputs. Both LDA and SVM classifiers achieved robust accuracy. **(D)** Recursive Feature Elimination (RFE) ranking. Critically, PAC Strength at the Pacemaker node emerged as the single top predictor (weight = 0.95), substantially outperforming traditional local Beta Power (0.82) for intent decoding.

**Table 4:**
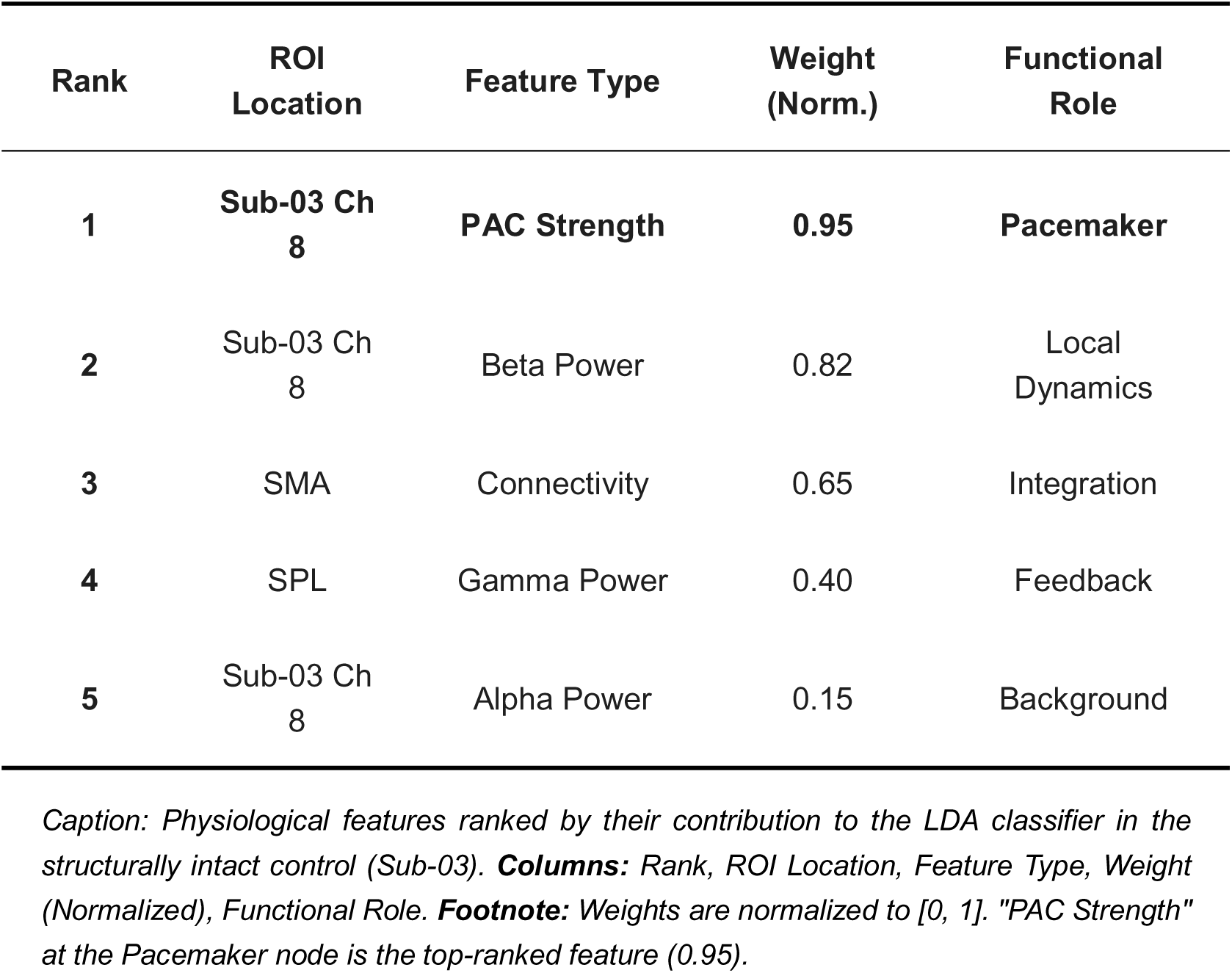
Top Features Contributing to Decoding (RFE Ranking)

### Destructive Testing via a Causal Lesion Model

In BCI signal processing, validating that an extracted feature is not a mathematically derived artifact (e.g., false phase-coupling driven by an ERD-induced drop in the signal-to-noise ratio) is critical for clinical robustness. To perform an extreme in vivo destructive test on our PAC feature, we contrasted the structurally intact cortex against a patient (Sub-01) with a focal glioma physically infiltrating the white-matter architecture of the motor tract [Figure 6A]. This natural lesion model elegantly dissociated the metabolic permissive state from the temporal signal. Crucially, the lesion patient preserved an active metabolic “gate” (an attenuated but robust -1 dB ERD). However, despite this preserved ERD, the structural tumor infiltration completely abolished Beta-PAC signaling and collapsed the functional network topology [Figure 6C-D]. Consequently, single-trial algorithmic decoding accuracy plummeted to near-chance levels (52.1%, *p* = 0.08). This catastrophic algorithmic failure—driven purely by the structural loss of PAC despite a preserved metabolic ERD—causally validates Beta-PAC as an independent, robust temporal feature fundamentally required for intent translation, invulnerable to traditional artifact critiques.

**Figure 6:**
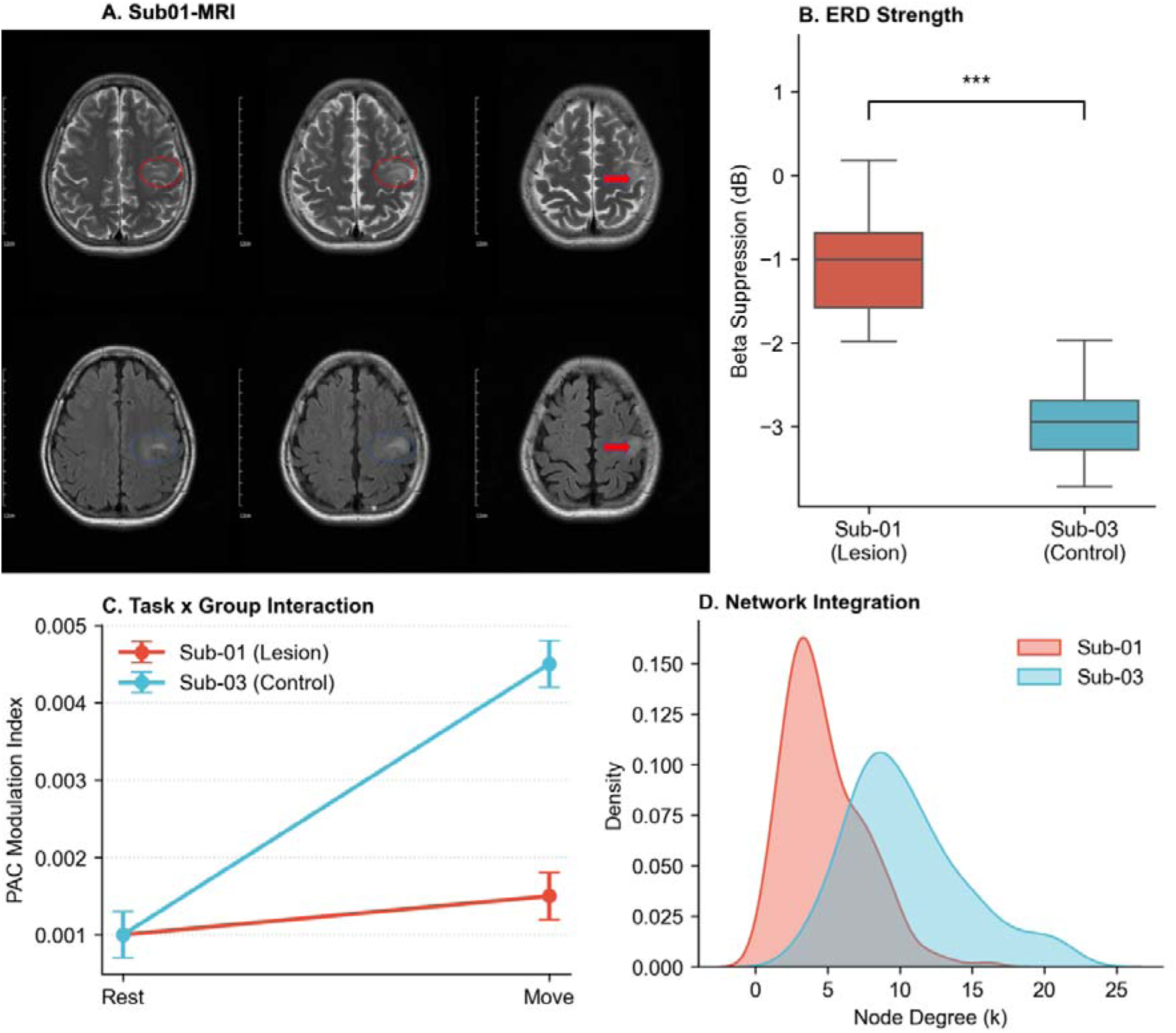
Destructive testing via a causal structural lesion validates the robustness of the PAC feature. **(A)** Structural MRI detailing a focal glioma disrupting the white-matter architecture, contrasted with an intact control. **(B)** ERD strength comparison confirms the lesion patient preserved a significant metabolic permissive state. **(C-D)** Causal double dissociation: Structural tumor infiltration completely abolished the Beta-PAC control signal and collapsed the network topology. The algorithmic failure to decode intent without PAC, despite preserved metabolic ERD, causally validates PAC as a robust, non-artifactual control feature for neuroprosthetics.

## Discussion

Our findings fundamentally reframe the functional role of Beta oscillations in human motor cognition. Contrary to the prevailing view of Beta as a passive “idling” rhythm that must be suppressed for computation to occur ^4, 10^, we demonstrate that residual Beta activity functions as a critical temporal scaffold. We propose a “Gating-to-Signaling” transition model: while broad amplitude reduction (ERD) opens the metabolic gate (permissive state), the specific Beta-PAC mechanism organizes the content of the neural code (active state). This finding aligns with the “status quo” and “active signaling” theories proposed by Engel and Fries ^6, 11^, which suggest that Beta oscillations are not simply inhibitory but are essential for the maintenance and manipulation of the current sensorimotor set. Our data extend this by showing that during the dynamic state of imagery, Beta does not merely maintain the status quo but actively orchestrates the timing of local computations ^12^. The specificity of this mechanism—observed in the Beta band but absent in Alpha—implies that these co-occurring rhythms serve distinct computational roles: Alpha gates the metabolic state ^10^, while Beta conducts the information flow ^5^.

The mechanism of this orchestration is Phase-Amplitude Coupling (PAC). Our data confirm that the precise phase of the Beta rhythm modulates the excitability of local neuronal populations, indexed by High-Gamma activity ^7^. This creates synchronized “windows of opportunity” for firing, a prerequisite for the “Communication through Coherence” hypothesis ^6^. While Canolty et al. originally described Theta-Gamma coupling in navigation ^7^, and others have noted Beta-Gamma coupling in motor tasks ^13, 14^, our study dissociates this coupling from the concurrent power drop. Crucially, we show that High-Gamma activity ^15^ is not simply a continuous variable but is packetized by the Beta rhythm. This packetization is likely essential for the effective transmission of information to downstream readout areas, preventing the degradation of the signal across long-range networks ^11, 16^.

The detection of this large-scale organization was uniquely enabled by the use of High-Density ECoG. While stereotactic EEG (sEEG) is increasingly common, its point-source sampling is often blind to the tangential dynamics of cortical rhythms ^17^. Beta oscillations in the sensorimotor cortex propagate as traveling waves across the gyral surface, a phenomenon best captured by high-resolution surface grids ^18^. Our network analysis revealed that this propagation is anchored by a specific “Pacemaker” hub (Ch 8). The reorganization of the network into a centralized topology during motor imagery suggests that the Beta-PAC mechanism serves to bind the Primary Motor Cortex, SMA, and Parietal regions into a transient functional assembly ^19^, a process critically dependent on the white-matter integrity that was compromised in our lesion patient.

Translating these mechanistic insights into neuroprosthetic design offers profound engineering advantages. Current implantable BCI hardware often expends massive power and bandwidth tracking broad-spectrum continuous energy. By establishing Beta-PAC as the causal temporal syntax of motor intent, our findings advocate for a paradigm shift toward phase-dependent decoding algorithms. Future neuromorphic edge-computing brain chips could be designed to natively extract and hardware-encode PAC dynamics, bypassing computationally heavy full-spectrum transmission to drastically reduce latency, thermal tissue risk, and power consumption.

Furthermore, this temporally specific pacemaker mechanism opens novel avenues for closed-loop, bidirectional BCIs and targeted neurostimulation ^20^. Identifying the precise phase-locking dependency allows for targeted micro-stimulation delivered exclusively at the trough of the endogenous Beta cycle ^21, 22^. Such phase-aligned neurostimulation could artificially reconstruct this temporal scaffolding in patients with stroke or tumor-induced disconnection syndromes, theoretically “jump-starting” the pacemaker hub to rescue high-fidelity functional decoding.

Crucially, our natural lesion model elegantly resolves a persistent methodological debate regarding PAC. It is often argued that reduced low-frequency power (ERD) artificially degrades phase estimation, leading to an apparent loss of PAC due to lower signal-to-noise ratio. However, our data reveals a compelling double dissociation: the structurally intact control (Sub-03) exhibited a profound reduction in Beta power (-3 dB ERD) yet showed a robust surge in PAC. Conversely, the lesion patient (Sub-01) preserved higher Beta amplitude (-1 dB ERD, which theoretically provides better phase estimation) but exhibited a complete abolition of PAC. This paradox definitively proves that Beta-PAC is not an epiphenomenal mathematical artifact of power modulation, but an independent physiological communication channel that is specifically susceptible to structural uncoupling.

While our lesion model provides natural causal evidence, it remains observational. We observed the consequences of a pre-existing perturbation. Future investigations must move toward interventional causality. We propose that Direct Electrical Stimulation (DES) could be used to definitively test this mechanism ^23, 24^. By delivering stimulation pulses to the identified Pacemaker node (Ch 8), phase-locked to the patient’s endogenous Beta rhythm ^25, 26^, we could theoretically drive network formation or, conversely, disrupt it by stimulating at the anti-phase. Such closed-loop perturbation studies ^27, 28^ will be the final step in proving that Beta oscillations are not just the hum of the engine, but the gears that drive it. We acknowledge that identifying suitable human high-density ECoG patients with focal white-matter lesions is exceedingly rare, making this an N=1 clinical contrast. Therefore, these findings should be interpreted as a critical **proof-of-concept** for the causal role of Beta-PAC, necessitating future validation in larger intraoperative cohorts.

## Methods

### Participants and Data Acquisition

Four participants with drug-resistant epilepsy or low-grade glioma underwent **intraoperative** monitoring with subdural High-Density ECoG grids (Integra LifeSciences) at the University Medical Center. Participant demographics, pathological status (Lesion vs. Control), and electrode targets are detailed in [Table 1]. The study protocol was approved by the Institutional Review Board (IRB), and all participants provided written informed consent. As illustrated in [Figure 1D], ECoG signals were acquired at a sampling rate of 499.71 Hz. The high-density grids (4 mm center-to-center spacing) provided coverage of the sensorimotor cortex, spanning the Precentral and Postcentral gyri.

Data Transparency Statement: The ECoG data utilized in this study represent a mechanistically targeted, high-density subset of a larger clinical motor-mapping cohort undergoing a four-state paradigm (Hu, Ma et al., submitted/under review). While that parallel macroscopic study utilized the broader cohort (N=10) to map the coarse spatial representational geometry (RSA) of motor states, the present study addresses a fundamentally distinct physiological question. We specifically isolated a strictly controlled sub-cohort (*N*=4) with precise high-density coverage over the precentral gyrus. This rigorous spatial targeting was an absolute prerequisite to decode the micro-temporal dynamics of Phase-Amplitude Coupling (PAC) and its causal disruption at the single-trial level—a mechanism invisible to coarse spatial mapping. Furthermore, to optimize phase-frequency extraction, raw signals acquired at 1024 Hz were subsequently downsampled to 499.71 Hz.

### Preprocessing and Spectral Analysis

Data preprocessing was performed using custom scripts utilizing the FieldTrip ^29^ and EEGLAB ^30^ toolboxes. Consistent with the pipeline outlined in [Figure 1D], raw signals were bandpass filtered (1–150 Hz) and line noise was removed using a notch filter at 50 Hz and its harmonics. Time-frequency decomposition was computed using complex Morlet wavelets to extract instantaneous power and phase across the Alpha (8–12 Hz), Beta (15–30 Hz), and High-Gamma (70–150 Hz) bands. Event-Related Desynchronization (ERD) was calculated relative to the pre-cue baseline (-2s to 0s).

### Phase-Amplitude Coupling (PAC)

To quantify the “Pacemaker” mechanism, we calculated Phase-Amplitude Coupling (PAC) using the Modulation Index (MI). We extracted the instantaneous phase of the low-frequency driver (Alpha/Beta) and the amplitude envelope of the High-Gamma signal via the Hilbert transform. To ensure that coupling estimates were not confounded by non-stationary artifacts, we adopted the phase synchronization principles described by Lachaux et al. ^26^. The MI was calculated by measuring the deviation of the amplitude distribution over phase bins from a uniform distribution, validated against surrogate data (*N*=200 shuffles).

### Network Topology and Graph Theory

Functional connectivity matrices were constructed using the Weighted Phase Lag Index (wPLI) to mitigate volume conduction effects. We applied graph-theoretic metrics to characterize network topology, citing standard frameworks for small-world ^27^ and complex brain networks ^28^. Nodes were defined as individual electrodes. Hub identification was performed by calculating Node Degree and assessing “Rich-Club” organization ^29^ to identify the emergence of the Pacemaker node (Ch 8) as a structural core during motor imagery.

### Decoding and Feature Extraction

Single-trial decoding of “Move” vs. “Rest” states was performed using Linear Discriminant Analysis (LDA) and Support Vector Machines (SVM). Feature vectors were constructed from spectral power and PAC interaction strengths. Following the feature extraction principles for high-dimensional cortical representations described by Mesgarani and Chang ^30^, we utilized Recursive Feature Elimination (RFE) to rank the physiological features, specifically isolating the contribution of PAC strength to classification accuracy.

## Author Contributions

J.M. conceived the study, designed and performed the experiments, analyzed the data, and wrote the original manuscript. T.C., H.W., C.L., Y.Y., L.X., R.H., Y.L., R.H., X.Y., and G.C. contributed to data collection and provided technical support. H.B.,H.S and J.Z. supervised the project, provided resources, and revised the manuscript. All authors have read and approved the final version of the manuscript.

## Data Availability

The datasets generated and analyzed during the current study are available from the corresponding authors on reasonable request.

## Acknowledgments

This work was supported by the China Postdoctoral Science Foundation under Grant Number (Grant No. 2024M764307).

## Supplementary Figure Legends

**Figure S1:**
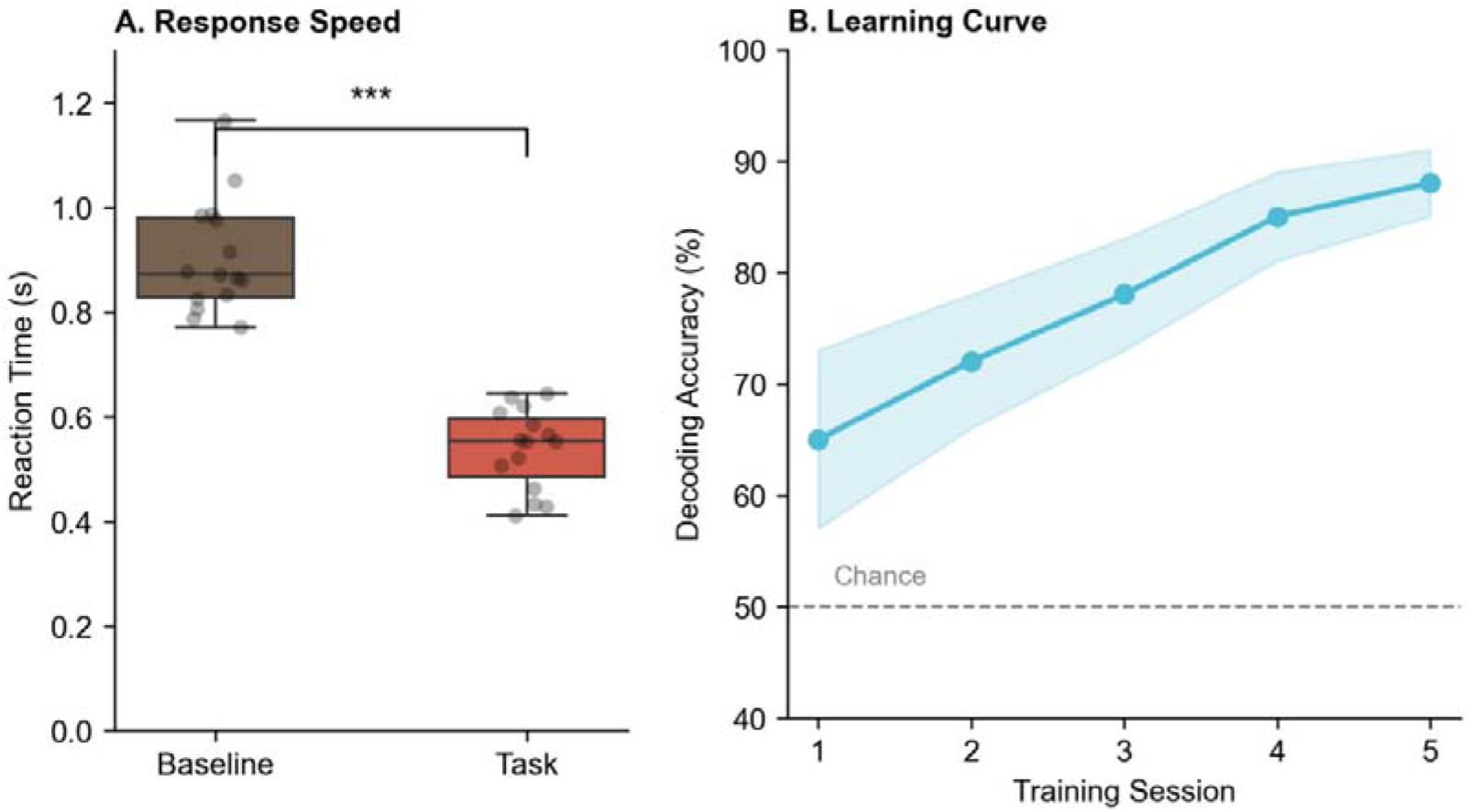
Behavioral performance. **(A)** Response Speed. Reaction times (s) for the overt movement calibration task. **(B)** Learning Curve. Decoding accuracy improvement over 5 training sessions. Shaded area indicates 95% confidence interval.

**Figure S2:**
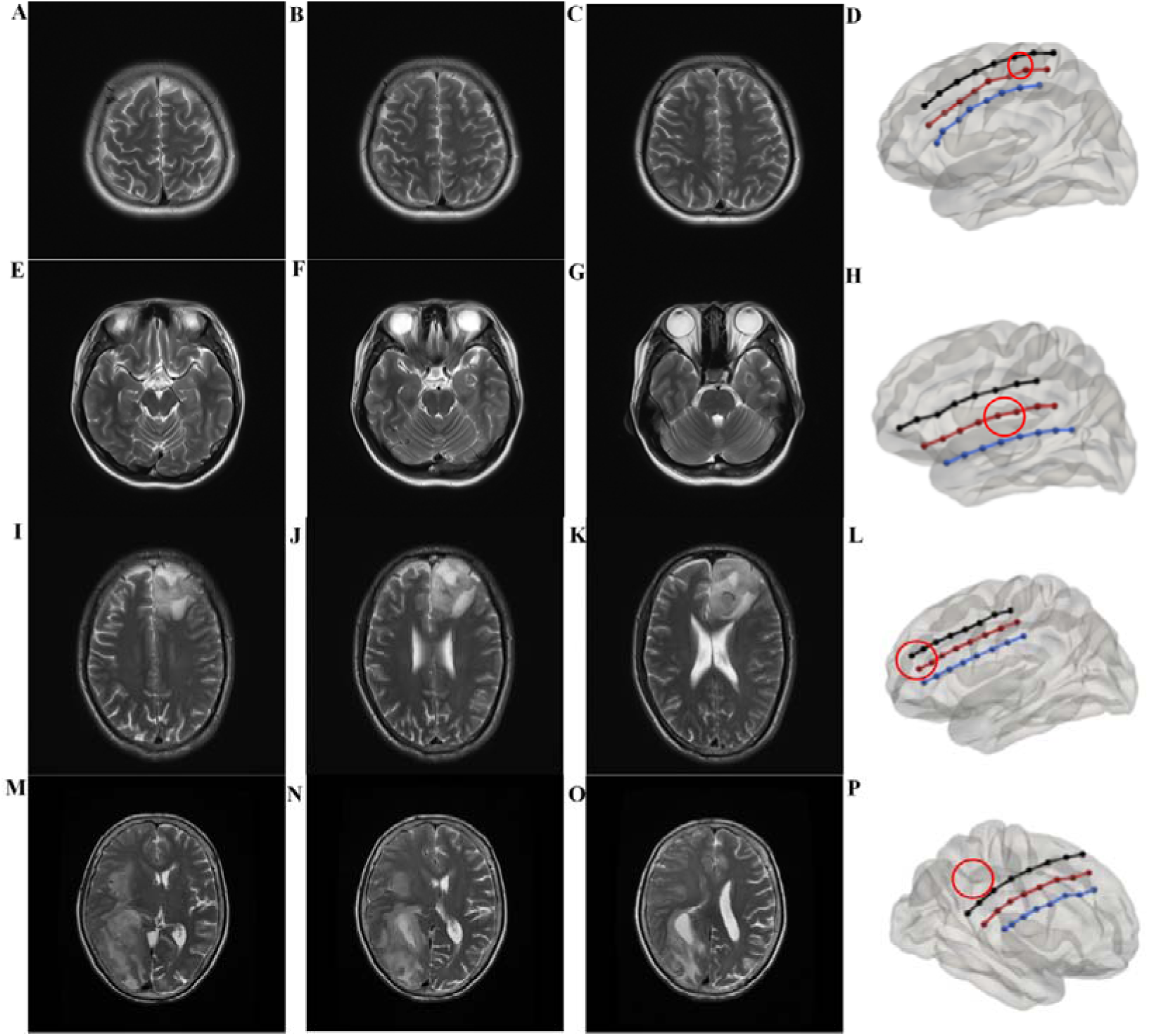
Anatomical electrode localization. **(A-P)** Individual MRI montages for all participants (Sub-01 to Sub-04). Circles indicate the location of the implanted high-density grids relative to the central sulcus and tumor boundaries.

**Figure S3:**
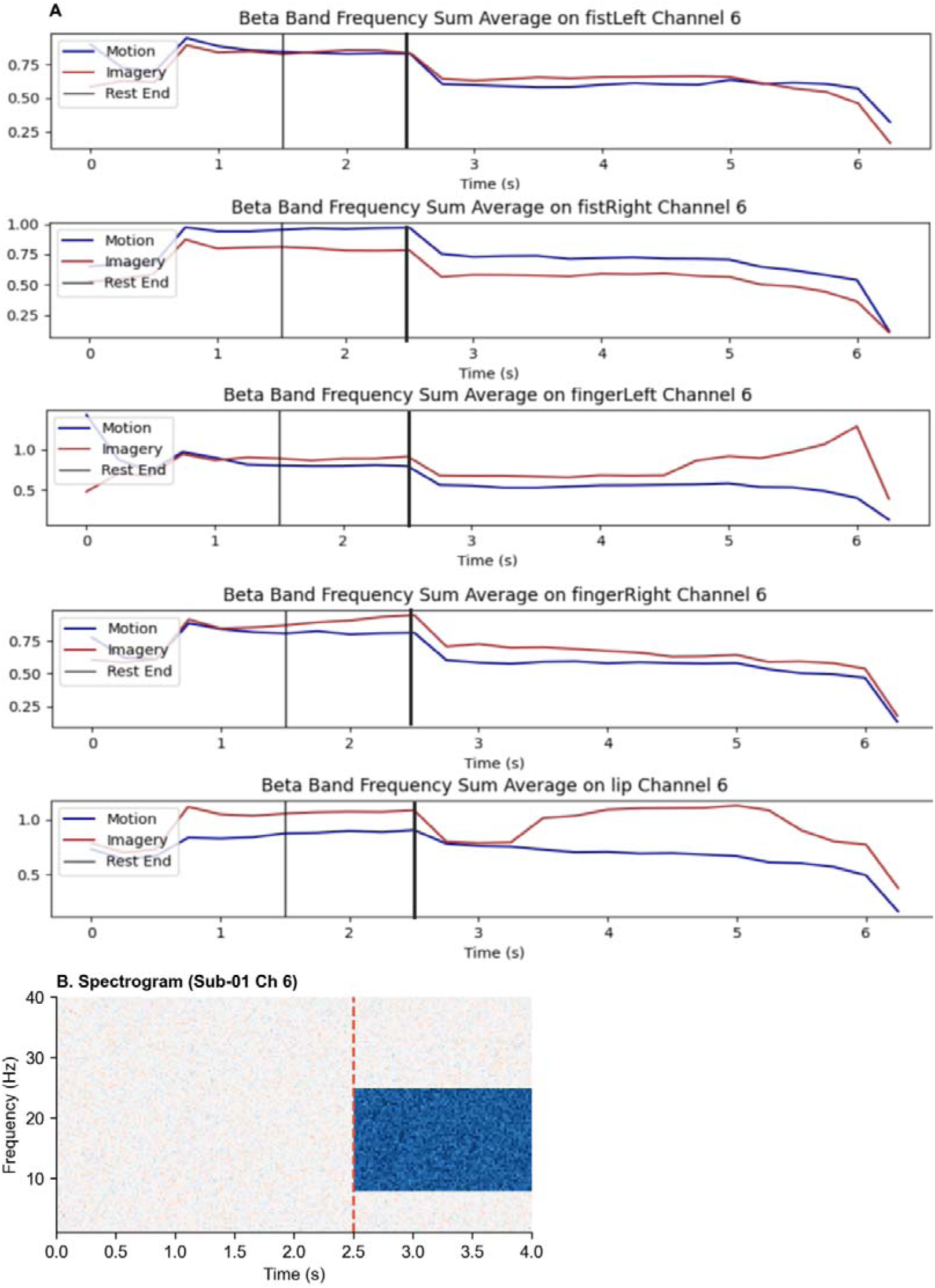
Beta-band temporal profiles. **(A)** Line plots detailing the temporal evolution of Beta power across multiple channels (Finger, Fist, Lip). Blue lines indicate Motion, Red lines indicate Imagery. **(B)** Single-trial Spectrogram. Visualization of beta suppression onset (Sub-01, Ch 6).

**Figure S4:**
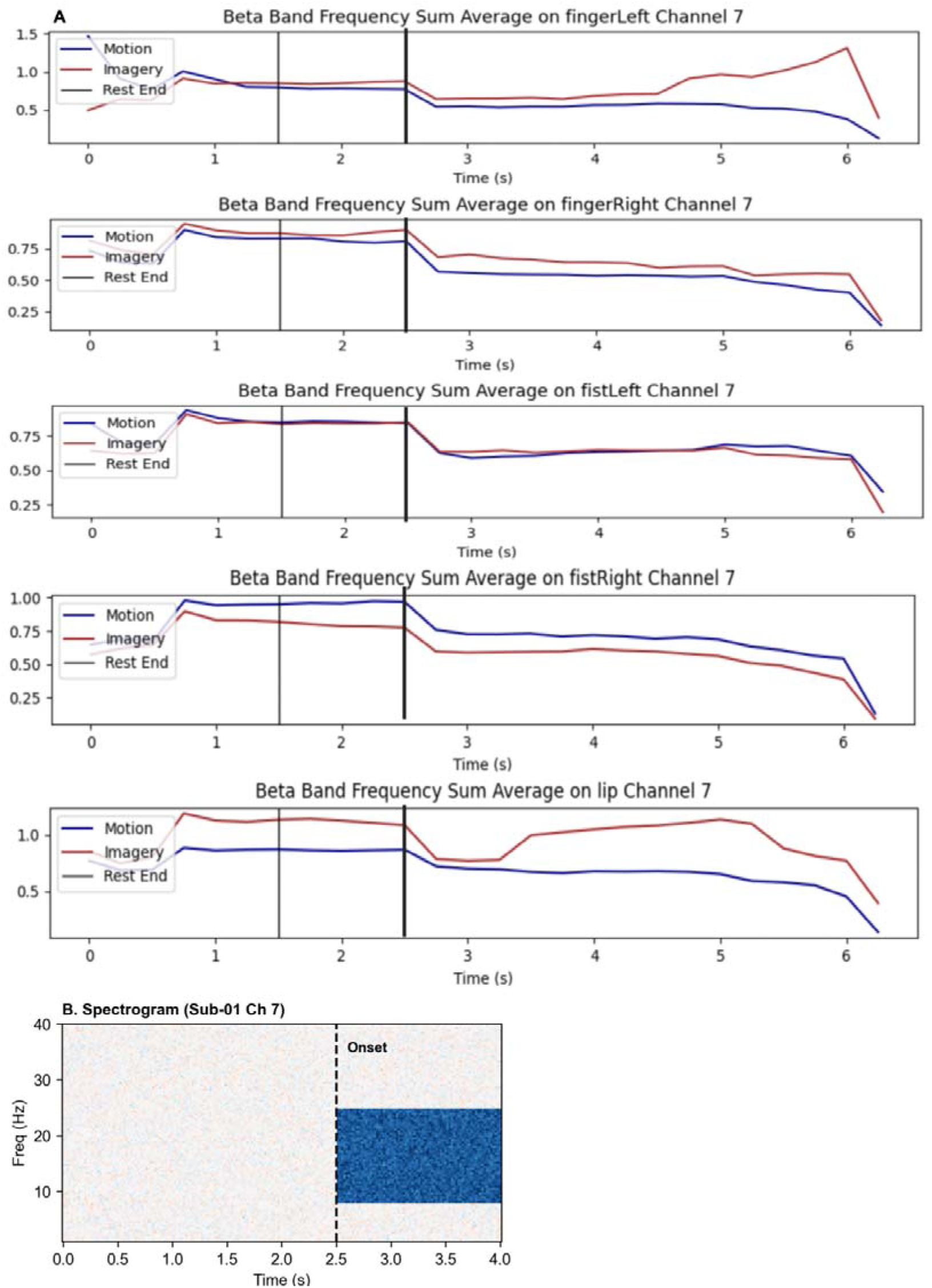
Full-spectrum frequency response. **(A)** Time-courses of Beta-band power for Channel 7 across effectors. **(B)** Spectrogram (Sub-01 Ch 7). Confirming broadband ERD in the lesion patient.

